# Age associated microbiome modulation and its association with systemic inflammation in a Rhesus Macaque Model

**DOI:** 10.1101/813667

**Authors:** Suresh Pallikkuth, Kyle Russell, Tirupataiah Sirupangi, Roberto Mendez, Daniel Kvistad, Rajendra Pahwa, Francois Villinger, Santanu Banerjee, Savita Pahwa

## Abstract

**Background:** As the individual ages, the immune system decreases in activity while chronic systemic inflammation increases. The microbiome is also affected by age, decreasing in beneficial microbes while increasing in pathogenic, inflammation inducing microbes with corresponding changes in their metabolic profile. While aging is known to affect both, links between the two have been hard to uncover.

**Methods:** Four young (age 3-6 years) and 12 old (age>18 years) Rhesus macaques were recruited for the study. PBMCs and plasma were collected to investigate immune cell subsets by flow cytometry and plasma cytokines by bead based multiplex cytokine analysis respectively. Stool samples were collected by ileal loop for microbiome analysis by shotgun metagenomics and serum, gut microbial lysate and microbe-free fecal extract was used to determine metabolomics by mass-spectrometry.

**Results:** Our aging NHP model recaptured many of the features of the age-associated immune alterations, with increased inflammation and alterations in immune cells subsets with lower number of CD4 T cells and a trend of age associated alterations in maturation subsets in older animals with lower naïve and higher central memory CD4 T cells. Older animals showed a significantly different microbiome from young animals with lower abundance of Firmicutes and higher Archaeal and Proteobacterial species. Correlation analysis showed a link between microbes in older animals with systemic inflammation. Analysis of metabolites in the serum and feces showed significant differences between specific metabolites between young and older animals that can influence age-associated morbidities.

**Conclusion:** These data support the age associated alterations in microbiome profile and its association with persistent systemic inflammation and metabolome changes. Further mechanistic studies are needed to understand the relationship between inflammation and microbiome. Nevertheless, this NHP model recapitulates human age associated changes in immune, inflammatory and microbiome profiles and can be useful for designing future studies targeting microbiome modulations in aging.

## Introduction

The world’s population is aging as every country in the world is experiencing growth in the number and proportion of older persons in their population. In 2010, an estimated 524 million people were aged 65 or older (8% of the world’s population) and by 2050, it is expected that about 1.5 billion (16% of the world’s population) people are aged 65 yrs or more. One of the complications of aging is its deleterious impact on the immune system leading to immunosenescence. Immunosenescence is a series of age-related changes that adversely affect the immune system and, with time, lead to increased vulnerability to various infectious diseases and comorbidities. The interplay between the age-related changes that affect different components of the immune system remains incompletely elucidated, and there is no clear understanding of which changes are primary, arising as a consequence of aging, and which might be secondary, adaptive or compensatory to the primary changes. Gaining knowledge of these mechanisms and interactions with multiple components of immune homeostasis will be critical for delaying the age-related decrease in immunity or preventing its consequences (1–5).

The process of aging involves a combination of alterations in the cells of the immune system, the microenvironment in lymphoid organs and non-lymphoid tissues and the circulating factors that interact with both immune cells and their microenvironment to insure the proper initiation, maintenance and cessation of immune responses as well as homeostasis of the immune system. Changes in the immune system as a result of aging are well documented (1, 3, 4, 6). Decreases in thymic activity result in less naïve cells being produced, lowering the proportion of naïve cells in both blood and secondary lymphoid organs (7). That decrease is accompanied by a relative increase in various populations of memory T cells (in particular, effector memory T cells. Antibody response to vaccinations has been shown to be reduced with age (1, 8). Additionally, inflammatory cytokines in circulation increase with age, leading to chronic systemic inflammation that negatively affects the host’s ability to mount immune responses (1, 9, 10). Age-associated chronic inflammation also negatively affects organ function, such as in the gut where inflammation leads to increased gut permeability, or “leaky” gut (2, 11, 12).

An increasing area of interest in immunology in the recent years is the impact of gut microbiome on regulating the immune homeostasis. The microbiome is the collection of microbes that exist in the gut of the host, and is known to undergo changes with age (2, 13–15). Early in adulthood, the microbiome is made up mainly by beneficial microbes from the Firmicutes and Bacteroides phyla (16). With age, the relative abundance of these phyla decrease as the microbiome becomes dominated by more pathogenic microbes from the Proteobacteria phylum. Changes in the microbiome that result in a more pathogenic state is termed microbial dysbiosis (2, 17–21). Age-associated inflammation is associated with microbial dysbiosis through a positive feedback loop where microbial products “leak” into the blood stream and increase inflammation (2, 17-19, 22, 23). In turn this inflammation causes the gut to be more permeable and allows more microbial products through, sustaining the feedback loop (2, 17–19, 22, 23). Aging research has experienced an extraordinary progress over recent years, particularly with the speculations that the process of aging could be controlled by maintaining the homeostasis of various genetic, biochemical and immunological processes. Many clinical issues, such as concomitant exposure to multiple drugs/antibiotics, dietary modifications and constipation, that generally accompany senescence are also closely correlated with perturbations in gut microbiome composition and functions. Gut microbiome is closely associated with several features of gut barrier integrity, intestinal pro- and anti-inflammatory balance, immune and cardio-metabolic health, and gut-brain axis. These evidences suggest that the gut microbiota may be associated with inflammaging and age-related chronic health conditions, and hence could be exploited as a putative target to ameliorate the aging process. While there is a known relationship between chronic systemic inflammation and microbial dysbiosis, elucidating specific mechanisms between the two has been challenging. To understand the relationship between aging, inflammation and microbiome, we conducted a study using aging non-human primates (NHP) model. Our aging NHP model recaptured many of the features of age-associated dysbiosis, inflammation and immune changes. We also found a relationship between age associated microbial dysbiosis with metabolic state of the host and microbiome.

## Results

### Higher systemic inflammation in older Rhesus Macaques

We enrolled 12 rhesus macaques with 8 old and 4 young animals in this study to investigate the effects of age on the immune system and microbiome. One of the hallmarks of aging is an increase in the systemic inflammation. We first wanted to test the systemic inflammation in this RM model by analyzing multiple plasma inflammatory cytokines (**Figure 1**). Old animals showed significantly higher plasma levels of acute inflammatory protein CRP **(Fig 1A)**, neopterin, a systemic marker of immune activation **(Fig 1B)**, and various inflammatory cytokines including TNF **(Fig 1C)**, IL-10 **(Fig 1D)**, IFNg **(Fig 1E)**, IL-8 **(Fig 1F**), IL-6 **(Fig 1G)**, and IL-2 **(Fig 1H**). Overall, these data support the presence of chronic persistent inflammation and immune activation in these aging NHP model.

**Figure 1:**
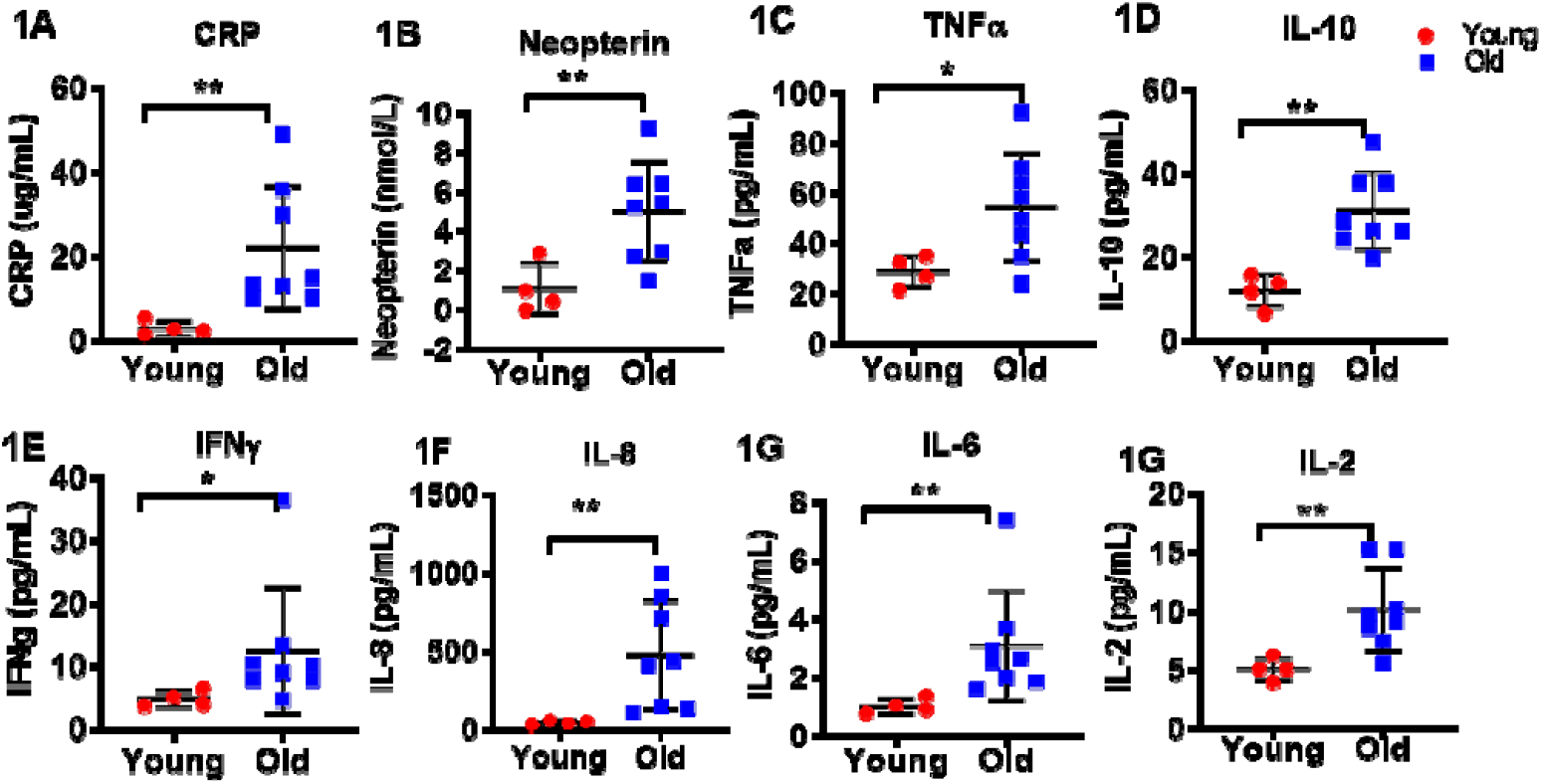
Plasma inflammatory cytokines in young and old Rhesus Macaques. ELISA kits were used to quantify levels of CRP (1A), and neopterin (1B) in young and old animals. Magpix multiplex analysis was used to quantify levels of TNFa (1C), IL-10 (1D), IFNg (1E), IL-8 (1F), IL-6 (1G), and IL-2 (1H) in young and old animals. Statistical analysis performed by student’s unpaired T test and significant differences with P<0.05 were identified by *.

### Age associated alterations in immune cell subsets are evident in these animals

We investigated the absolute numbers and frequencies of immune cells in circulation by analyzing the PBMC using flow cytometry (**Figure 2**). While CD4 and CD8 absolute counts are not given by total blood count test, we used the lymphocyte absolute counts with CD4 and CD8 frequencies from flow cytometry to calculate CD4 and CD8 absolute counts. A trend of lower absolute numbers of lymphocyte **(Fig 2A)** and a trend of higher numbers of monocytes **(Fig 2B)** were observed in older animals but the differences were not statistically significant. Within the lymphocytes, we found a significantly lower number of CD4 T cells **(Fig 2C)** in older animals while absolute number of CD8 T cells **(Fig 2D)** did not differ significantly between young and old animals.

**Figure 2:**
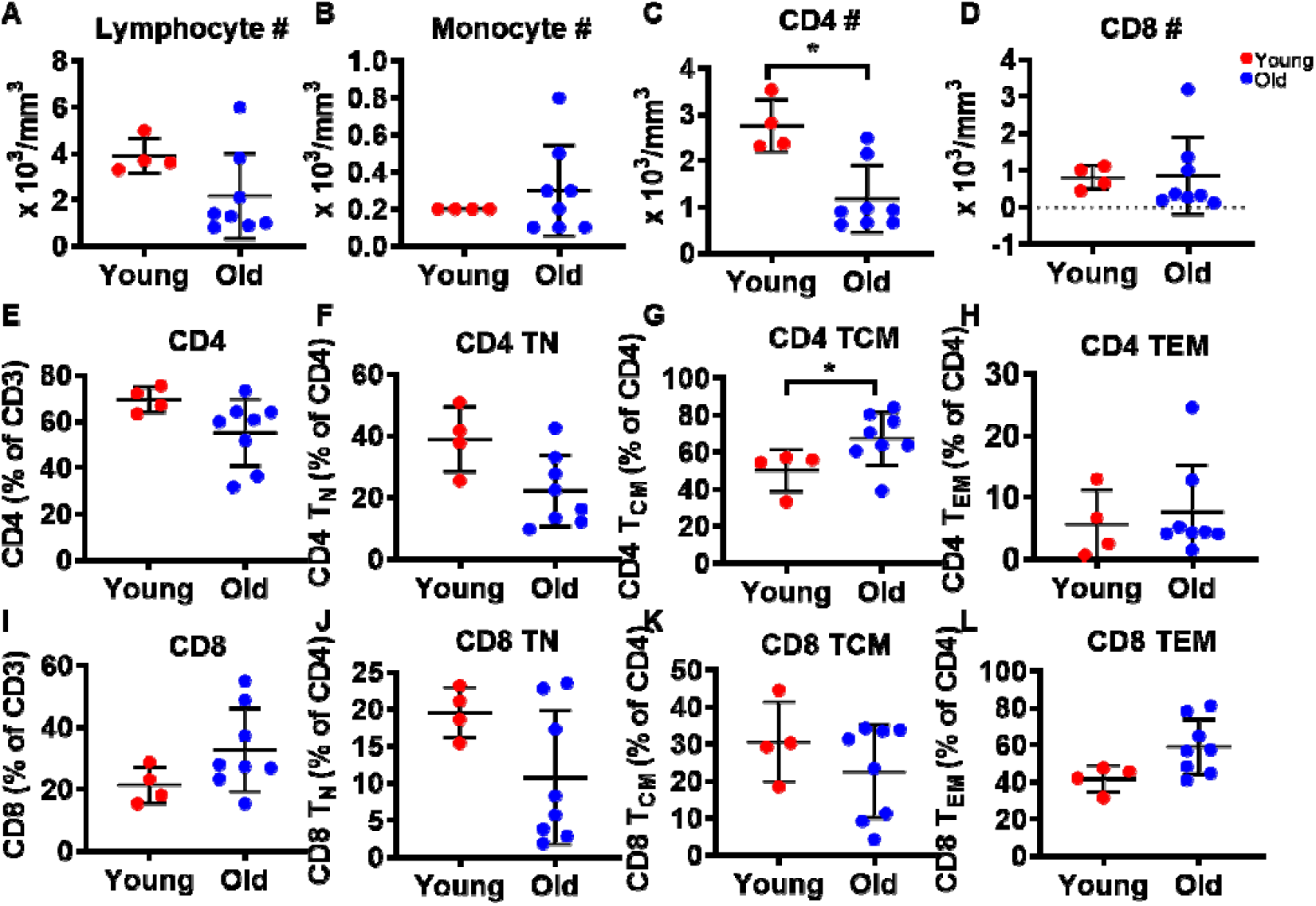
Absolute numbers and frequencies of CD4 and CD8 T cells in young and old Rhesus Macaques. Blood counts was done to determine absolute counts of and flow cytometry for frequencies of CD4 and CD8 T cells from PBMC. Scatter plot showing absolute numbers of A), lymphocytes, B), monocytes, C), CD4 and D), CD8 T cells in young and old Rhesus Macaques. Scatter plot showing frequencies of E), CD4, F), naïve CD4 (T_N_) F), central memory (T_CM_) and G), effector memory (T_EM_). H), CD8 T cells I), naïve CD8, J), TCM CD8 and K), TEM CD8 T cells. Statistical analysis performed by student’s unpaired T test and * indicates p value <0.05.

We next analyzed the frequencies of CD4 and CD8 T cells and their maturation subsets between the young an old animals. In agreement with lower numbers of CD4 T cells, we found a trend of lower frequencies of CD4 T cells (Fig 2E) in the older animals although the difference was not significant. Analysis of CD4 T cell maturation subsets revealed a trend of lower frequencies of naïve (defined as CD4^+^CD28^+^CD95^−^) (Fig 2F) along with a higher frequencies of central memory(defined as CD4^+^CD28^+^CD95^+^) CD4 T cells (Fig 2G) in the older animals while CD4 effector memory (defined as CD4^+^CD28^−^CD95^+^) subset (Fig 2H) did not differ between the study groups. While there were no significant differences in frequencies of CD8 cells (Fig 2I) or subsets, similar trends were observed. A trend of lower CD8 naïve (Fig 2J) and central memory (Fig 2K) frequencies were noted in the older animals while effector memory (Fig 2L) frequencies were higher in older animals. Taken together, our data support an age associated increase in inflammation and alterations in immune cell subsets in this aging NHP model that are in agreement with human studies.

### Microbiome between young and old animals show significant differences

We next analyzed the gut microbiome profile of these animals to investigate the age associated microbiome changes. To test the gut microbiome, we collected stool samples from young and old rhesus macaques then analyzed them by shotgun metagenomics sequencing (**Figure 3**, **4**). At first, we analyzed the diversity changes between the two groups (α-diversity) using three different indices, namely Shannon (**Fig 3A**), Chao1 (**Fig 3B**) and Simpson (**Fig 3C**). While there was a trend towards decrease in the Shannon index and increase in Chao1 and Simpson indices with age, none of the changes were statistically significant. Next, we analyzed the group-wise compositional differences using Bray-curtis matrix (β-diversity; Principal coordinate analysis or PCA). PCA analysis showed a clear difference in microbiome composition between young and old animals without any overlap, with significant differences between the two **(Fig 3D)**.

**Figure 3:**
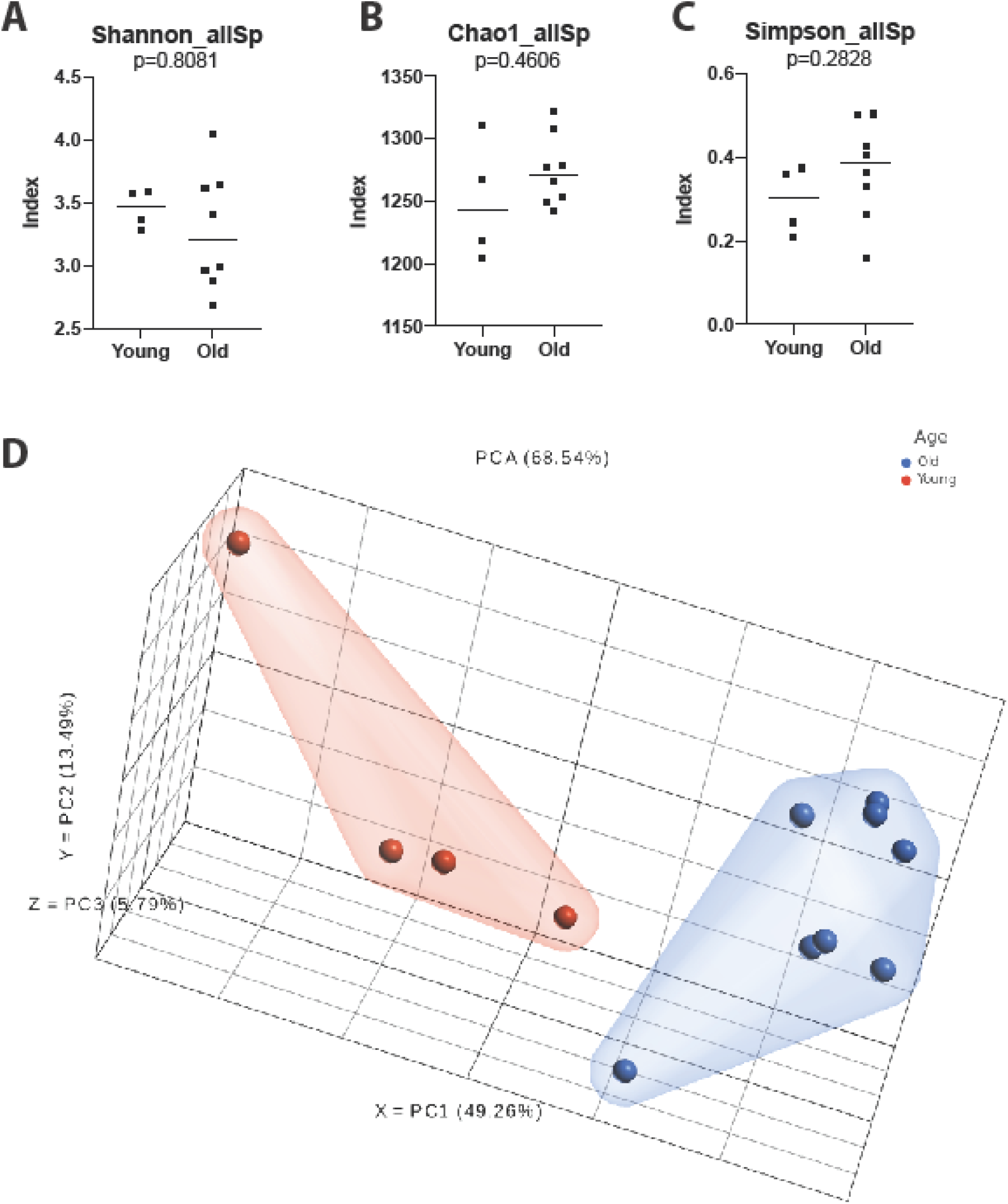
Gut microbial diversity indices of old and young animals. ◻-diversity indices, namely Shannon (A), Chao1 (B) and Simpson (C) do not exhibit any significant differences between old and young animals. Qualitative differences between the two age groups (◻-diversity), rendered as PCA plot (Bray-curtis) shows distinct clustering between the two. Test of significance (A, B and C)—Mann-Whitney U test, using Graphpad Prism. P-values are exact.

**Figure 4:**
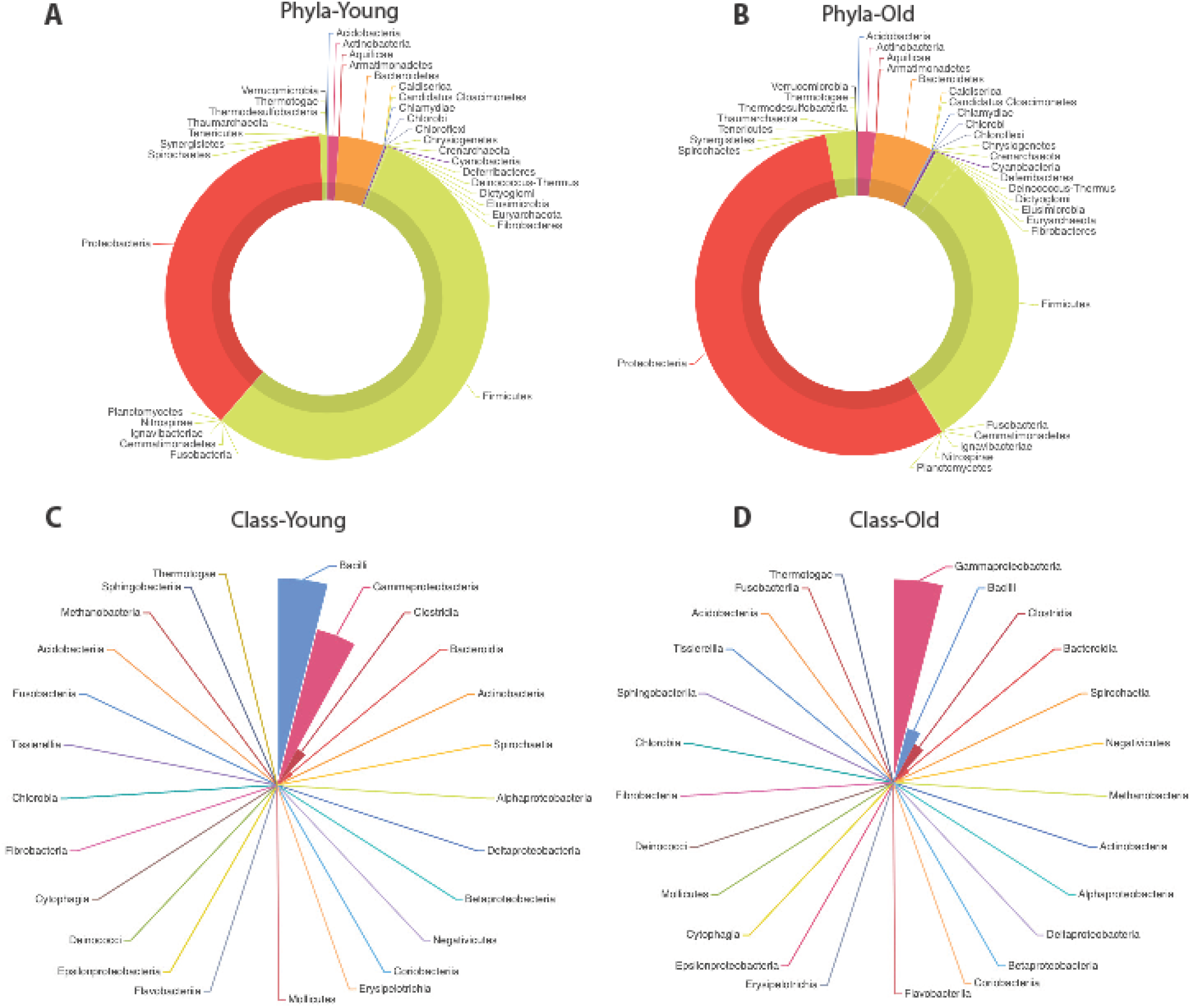
Phylum and Class-level differences with age. *Rhesus* gut microbiome is Proteobacteria and Firmicutes dominated (A, B), followed by phylum Bacteroidetes. In young animals (A), we saw a Firmicutes dominated microbiota that becomes Proteobacteria dominated with age (B). At the class level, Proteobacteria and Firmicutes dominates the landscape. In young animals (C), microbiome was Bacilli (Firmicutes) dominated, followed closely by Gammaproteobacteria (Proteobacteria). Clostridia (Firmicutes) and Bacteroidia (Bacteroidetes) were seen to be the other dominant classes in these animals. In older animals (D), in contrast, Gammaproteobacteria is seen to be the most dominant class with greatly diminished representation from Bacilli as the second most abundant class.

When we looked at the bacterial phyla and classes driving the differences between the microbiome in young and old animals, it was evident that gut microbiome in young old animals was dominated by Proteobacteria and Firmicutes (**Fig 4A, 4B**), followed by phylum Bacteroidetes. In young animals (**Fig 4A**), showed a Firmicutes dominated microbiota that becomes Proteobacteria dominated with age (**Fig 4B**). In older animals, in addition to expansion of Proteobacteria in relative abundance, we also see relative expansion of Bacteroidetes, Tenericutes and Verrucomicrobia, all at the expense of Firmicutes. Firmicutes, however, still remained the second largest phylum on old animals. At the class level (**Fig 4C, 4D**), classes within Proteobacteria and Firmicutes dominated the landscape. In young animals (**Fig 4C**), microbiome was Bacilli (Firmicutes) dominated, followed closely by Gammaproteobacteria (Proteobacteria). Clostridia (Firmicutes) and Bacteroidia (Bacteroidetes) were seen to be the other dominant classes in these animals. In older animals (**Fig 4D**), in contrast, Gammaproteobacteria is seen to be the most dominant class with greatly diminished representation from Bacilli as the second most abundant class. Clostridia and Bacteroidia occupy the third and fourth most dominant position within the old group and their relative abundance in unchanged compared to the microbiota in young animals.

At the species level, 55 species were significantly different between the young and old *Rhesus* gut microbiome (**Supplementary Figure 1**). As expected with minor exceptions, most of the species showing diminished relative abundance in old animals belong to phylum Firmicutes, while the ones with elevated relative abundance are members of phylum Proteobacteria.

### A potential association between systemic inflammation and gut microbiome in older animals

To better understand the relationship between the microbiome and host immunity, a correlation analysis between specific microbial species and plasma inflammatory cytokines was performed. The list of over 600 microbes identified during the microbiome was arranged according to relative abundance difference between young and old animals. Twenty-five microbes from each end of the list, representing microbes most abundant relatively in young or old animals, were then correlated with plasma cytokines. The results were graphed using a heat map (**Fig 5**), where red signifies positive correlations while blue signifies negative correlations and * indicates the significant correlations. Microbes more abundant in older animals were almost all positively correlated with plasma inflammatory cytokines with many microbes showed significant correlations with neopterin, CRP, TNFa, IL-2, IL-6, IL-8 and IFNg. Majority of the microbes correlated with neopterine and CRP. Microbes more abundant in younger animals showed a trend of negative correlation with the same cytokines, with neopterin showing the significant correlation.

**Figure 5:**
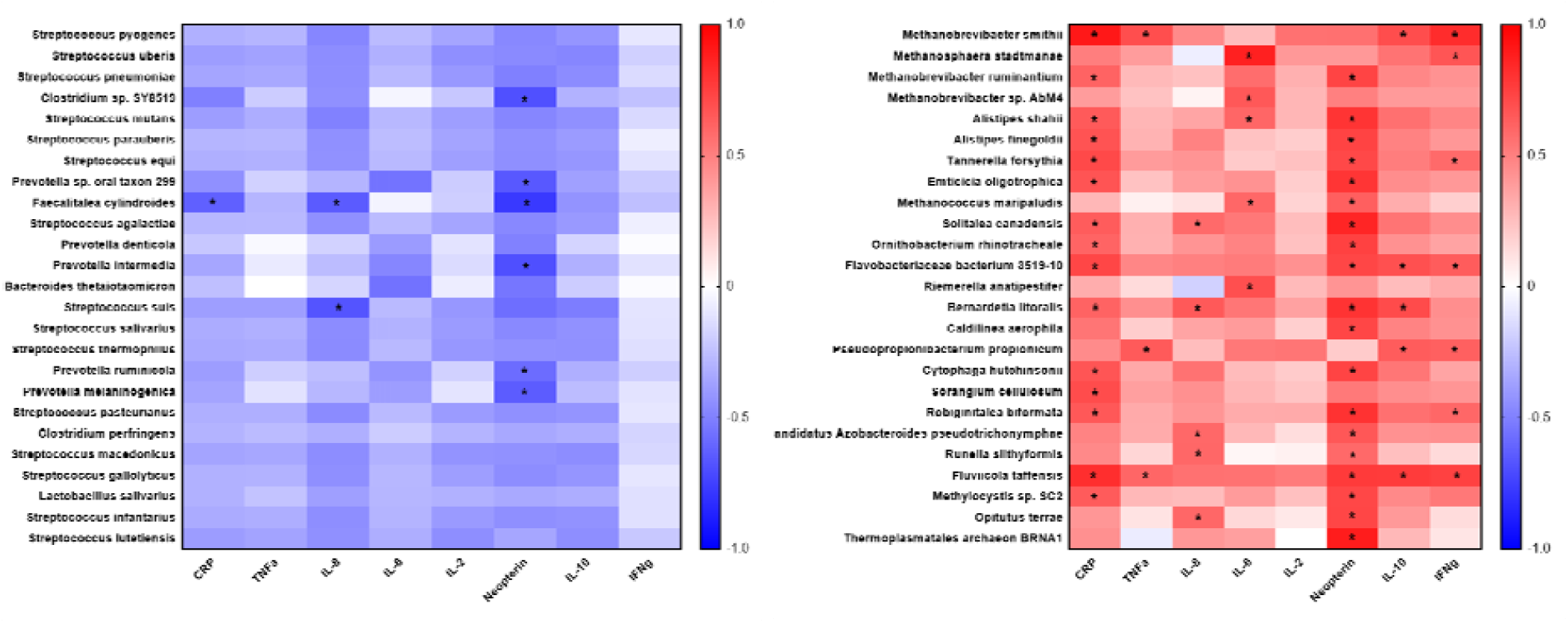
Correlation analysis of prevalent microbes in young or old Rhesus Macaques and plasma inflammatory cytokines. Microbes were selected based off increased prevalence in young (5A) or old (5B) rhesus macaques and then correlated with plasma cytokines. Positive correlations are in red. Negative correlations are in blue. Statistical differences are identified by *. Older animals showed significantly more correlations with plasma cytokines, all of which were positive correlations.

### Bacterial association networks significantly change with age

Bacterial association was calculated using Markov Clustering Algorithm (MCL), which evaluates correlations of vector abundance of bacterial species within each group. This results in clustering of species that are most likely to co-occur together in a network, using correlation values as the distance matrix. This also allows for each species to participate in multiple clusters. As shown in **Fig 6A**, microbiome of young animals exhibited a single large super-cluster at microbial homeostasis. However, with age, major constraints have been introduced into the network structure with the emergence of satellite clusters and scattering of the super cluster (**Fig 6B**). In old animals, various newly emerging breakout clusters could be seen with known bacterial composition (Representative clusters numbered, and composition shown in **Supplementary Figure 2**). It is conceivable that age-induced relative changes in the microbial composition would result in equitable changes in bacterial associations as well. For example, in cluster 1 in Fig 5B, the bacteria belong to phyla Archaea, Thermotogae, Proteobacteria and Bacteroidetes (Supp Fig 2). One of the common properties in all constituents of cluster 1 is that they are all extreme thermophiles. Similarly, in cluster 2, with the exception of *Lactobacillus casei*, all constituents are known pathobionts (i.e. potential mammalian pathogen). Hence, with age-induced microbial dysbiosis, constraints are introduced into the homeostatic microbial networks and bacteria of known traits seemingly cluster together, forming their own niche. This could also result in change in their metabolic profiles and overall metabolic profile of the microbial compartment.

**Figure 6:**
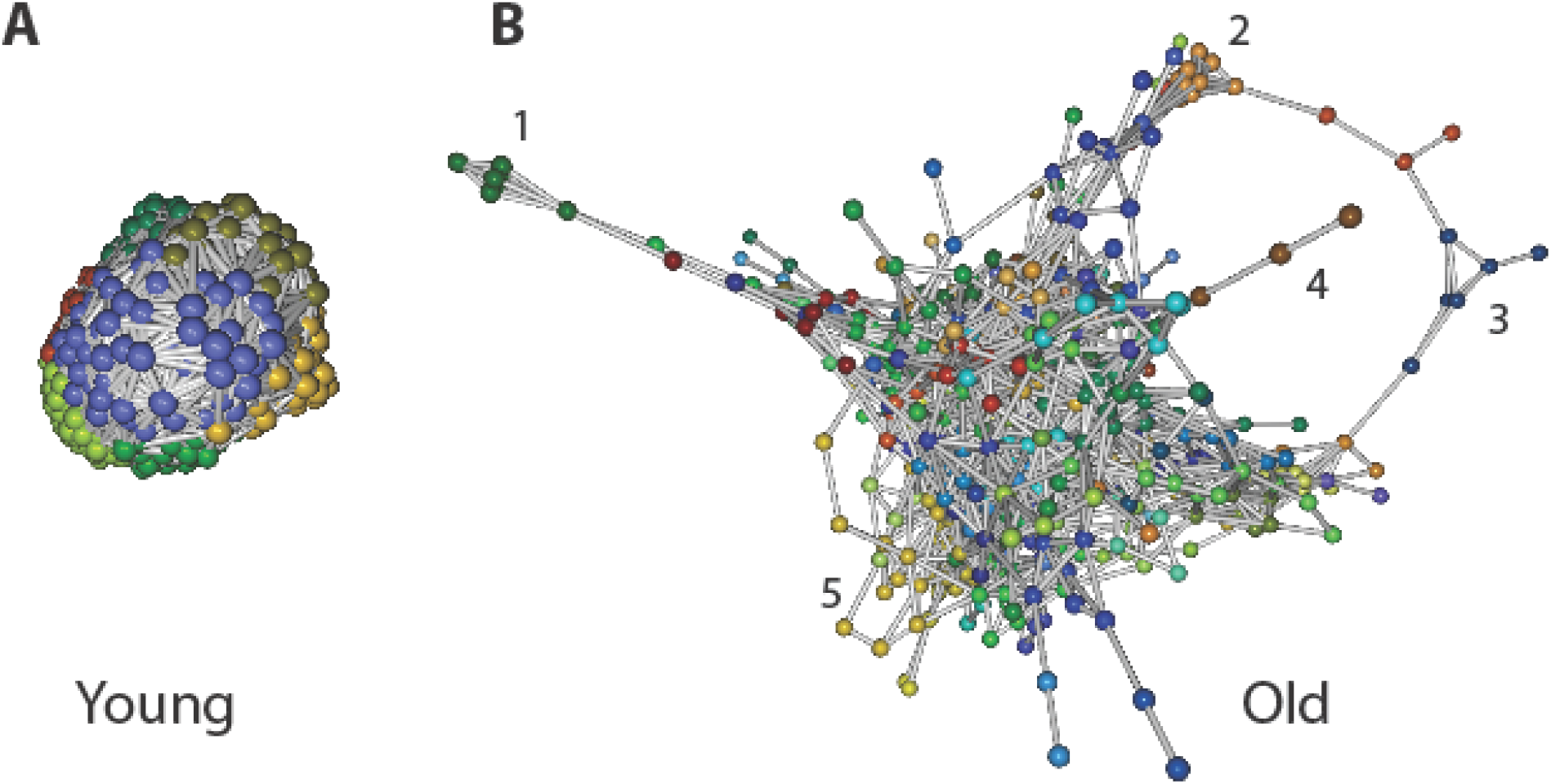
Bacterial association networks and differences with age. Markov Clustering Algorithm (MCL) evaluates correlations of vector abundance of bacterial species within each group, thereby generating a cluster map by using correlation scores as distance. Figure shows that microbiome of young animals group into a single large super-cluster at microbial homeostasis (A). However, with age, major constraints have been introduced into the network structure with the emergence of satellite clusters and scattering of the super cluster (B). Cluster identities are listed in Supplementary figure 2

### Serum metabolites show significant differences between young and old animals with strong correlation with altered microbial metabolism

One of the ways the microbiome can affect the host is through their own metabolism, producing specific metabolites in different abundances that make their way into the host circulation. Metabolites in the serum of young and old animals was analyzed by mass-spectrometry based Isotope Ratio Outlier Analysis (IROA^®^) as we have described before (24). Investigation in serum from young and old animals showed a few metabolites that were significantly different between the two groups (**Figure 7A**). Old animals showed significantly higher levels of hypoxanthine, L-arginine, and an unidentified compound with a chemical composition of C7H11N6O4. Hypoxanthine is used in purine metabolism to make nucleic acids while L-arginine is an amino acid used in the polyamine pathway to make nucleotides. Younger animals showed significantly higher levels of L-nicotinamine, fructoselysine, N-acetylneuraminate, leucyl phenylalanine, and 2’,3’-cyclic uridine monophosphate (UMP). L-nicotinamine is a plant-derived siderophore, used by bacteria and fungi for amino acid synthesis. Fructoselysine is a bacterial product, composed of fructose and lysine that is broken down for use as energy or amino acid production. N-acetylneuraminate is consumed by bacteria as a carbon source. Leucyl phenylalanine is used in amino acid production. Lastly 2’,3’-cyclic UMP is used in nucleic acid production in bacteria, and participates in mitochondrial ATP to ADP reduction in mammals. Upon mapping these metabolites on pathways (Old over Young), most of the biosynthetic pathways are seen to be upregulated on the old animals (**Figure 7B**). At the same time, except carbohydrate and secondary metabolite catabolism, all degradation pathways were seen to be higher in older animals (**Figure 7C**). The point that needs to be emphasized here is that observations in Figures 6B and C are based on presence/absence and relative abundance of metabolites from the serum of old and young macaques, which may or may not be a consequence of gene and pathway activation. However, downstream participation of these metabolites in various pathways would definitely be consequential to the host.

**Figure 7:**
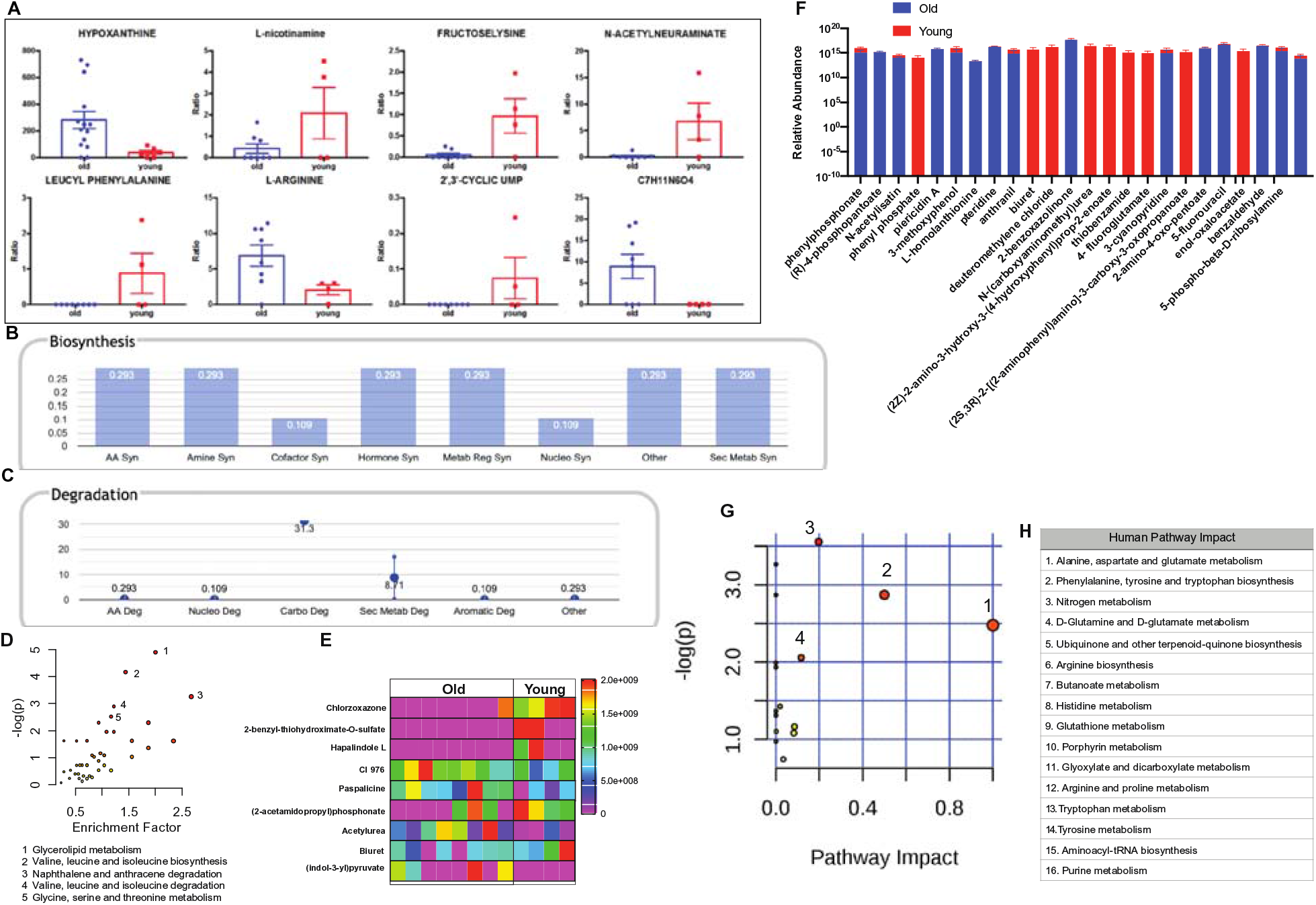
Crosstalk between the bacterial and host metabolic pathways. Mass-spectrometry based Isotope Ratio Outlier Analysis (IROA^®^) showed a few metabolites that were significantly different between the two groups (A). Upon mapping these metabolites on pathways, most of the biosynthetic and stress pathways are seen to be upregulated on the old animals (B). Additionally, with the exception of carbohydrate and secondary metabolite catabolism, all degradation pathways were seen to be higher in older animals (C). LC-MS/MS analysis of the pure microbial fraction and pathway enrichment analysis showed that glycerolipid, amino acid and secondary metabolite metabolism predominated the metabolic landscape of both groups (D). Upon mapping the significantly different metabolites between old and young macaques (E) to metabolic pathways, a synchrony was observed between microbial and host biosynthetic and degradation pathways. In the microbe-free fecal compartment, after adjusting for dietderived metabolites, several metabolites were significantly altered, that were mapped to stress pathways in the aged animals (F). When all significantly changing metabolites in old and young animals were mapped for human pathway impact (G; PathwayAnalyst 4.0),secondary metabolite synthesis, nucleotide synthesis and amino acid degradation were still at synchrony with bacterial and host metabolic status (H).

To understand the metabolic landscape of the microbial compartment from the two groups, we next isolated metabolites from the microbial and microbe-free fecal matter from the animals of the two groups. LC-MS/MS analysis of the pure microbial fraction and pathway enrichment analysis showed that glycerolipid, amino acid and secondary metabolite metabolism predominated the metabolic landscape of both groups (**Figure 7D**) with significantly different metabolites shown in **Figure 7E**. When we looked at the microbe-free fecal compartment (old over young), secondary metabolite synthesis, nucleotide synthesis and amino acid degradation were still at synchrony with bacterial and host metabolic status (**Figure 7F, 7G** and **7H**). Several metabolites were significantly different in old and young maquaques (**Figure 7F**), which were mapped to various human pathways using KEGG orthologies. *Macaque* chow was used as a control for this experiment and all metabolites present in the chow, including those common in bacterial fraction were not considered for the differential analysis. Significantly different metabolites were mapped on human pathway impact using MetaboAnalyst 4.0 and significantly impacted pathways were identified (**Figure 7G** and **7H**).

## Discussion

In this study, we investigated the association between age associated changes immune homeostasis and inflammatory microenvironment on gut microbiome composition and its impact on host metabolome using an aging NHP model. We were able to recapitulate age associated alterations in immune cell compartment and systemic inflammation in this animal model. We found significant increases in several cytokines associated with inflammation and soluble markers of immune activation in older animals. Increases in IL-6, IFNγ, and TNFα are expected as these are known inflammatory cytokines that act on a systemic level, promoting chronic inflammation that’s associated with age (25). Increased CRP and neopterin are indicative of an increased systemic immune activation inflammatory state in older animals (26, 27). Neopterin, a metabolite of GTP, is produced in macrophages upon stimulation by IFNg released by activated T cells indicating an ongoing immune system activation and inflammatory response in the older animals. Consistent with increase in neopterin, we also found an increase in IL-6 and IL-8 supporting a generalized monocyte/macrophage activation contributing to systemic inflammation in the older animals (28). While IL-10 is an anti-inflammatory cytokine, it can be rationalized that it’s elevated in older animals as an attempt by the immune system to dampen systemic inflammation and maintain immune regulation. Overall these results are still in line with what was hypothesized and are encouraging for further study.

Another significant change as the result of the aging process is the alterations in the immune cell subsets in the circulation. Decreases in frequencies of CD4 T cells was expected in older animals as they don’t produce as many new immune cells, which was also observed in the decreased frequency of CD4 naïve cells. As a result, there was a significant increase in the frequency of CD4 central memory in older animals indicating an ongoing T cell differentiation without production of newer cells. It is possible that ongoing inflammation and immune activation may support the terminal differentiation of immune cells in older animals. Despite increased frequencies of CD8 in older animals, there were decreased frequencies of CD8 naïve and CD8 central memory, with increased frequencies of CD8 effector memory. The observed decrease in the absolute numbers of lymphocytes in older animals was not surprising as age associated decrease in cell numbers have been reported. Interestingly absolute counts of monocytes and CD8 T cells increased with age, though not significantly. The only significant change was observed in the decrease of absolute count of CD4 T cells in older animals. Overall though there were significant changes in immune cell frequencies that were based on age.

Our most interesting data came from the microbiome. Significant differences were observed between the microbiome of young and old animals. Changes in the microbiome as a result of age are well known (2, 15, 19, 20, 23), with the microbiome of older animals showed a more proinflammatory profile. The significant differences in microbiome composition shown in the PCA plot was further supported by the class and species-level differences that further showed changes in the taxonomic composition of microbiome with aging in these animals. Unlike in humans, dietary influence on microbiome composition could be minimal in these animals as these animals were fed on the same diet, and observed differences in microbiome composition must be closely related to aging process. It is also important to note that even with a modest number of animals per group in this study, we were able to identify a significant alteration in microbiome profiles indicating the strong influence of aging on microbiome changes at individual animal level. In addition, we expanded our investigation of composition beyond diversity to evaluate clustering patterns, noting differences between the two age groups, with respect to pattern and clustering due to relative perturbations in bacterial populations. These changes in community behavior align with biological context, which demonstrate that compositional changes in microbiota due to disease is not a singular change. Changes in relative abundance of microbial species results in imposition of constraints on co-occurrence network(s), resulting in induction/expulsion of other species, or complete rearrangement of networks (29). As mentioned in the results, these changes are also associated with alteration of biochemical behavior of individual microbes, often resulting in enhanced expression of virulence factors and metabolites, that have influence on host immunity and inflammation with age.

The next step was looking into potential mechanisms linking the microbiome and host immune system and how age impacts these interactions. There’s plenty of evidence of cross-talk between the microbiome and immune system, however figuring out the specific mechanisms going on between the two have been difficult. To elucidate a potential mechanism, we first tried to identify potential associations between systemic markers of inflammation and immune activation and microbes. Our data showed that microbes that are more prevalent in young animals showed mainly negative correlations with systemic inflammation while microbes more prevalent in older animals showed mainly positive correlations with plasma multiple markers of inflammation and immune activation. Specifically, many predominant microbes in older animals correlate with CRP and neopterin, which again are generally markers of systemic immune activation and inflammation. It’s known that with age the permeability of the gut increases, allowing more bacterial products to translocate to the blood and promote inflammation (2, 11, 12, 30). Different bacterial products from different microbes could elicit stronger inflammatory responses in the blood, feeding back onto the gut to promote gut permeability. Through this way the microbes in the gut and the host inflammatory response create a positive feedback loop that accumulates with age. Considering the microbes in older animals showed more significant correlations, investigating these specific microbes and their potential for activating the immune system could help shed light on how the microbiome affects systemic inflammation.

The other method done to investigate future directions was analyzing the available metabolites of the animals. The microbiome is known to play a big role in metabolism, breaking down food and producing a number of different metabolites that are absorbed by the host. In turn these metabolites can have a number of different effects on the host body, including affecting and altering the immune system. Thus through metabolism the microbiome can have an indirect effect on host immunity. Figure 6 shows the significant differences in metabolites seen in the serum of young and old animals. One of the compounds, C7H11N6O4, were described structurally but not identifiable. Hypoxanthine, increased in older animals, is oxidized to uric acid at the end of purine metabolism, releasing reactive oxygen species (ROS) (31). 2,3-cUMP, which is abolished in the old animals, is a predominant bacterial product, that can be converted to nucleoside-3-phosphate in eukaryotes and it’s inhibition has be shown to induce mitochondrial stress by modulating Akt and GSK3b activity (PMID: 30405014). Fructoselysine is an Amadori product that can be broken down by commensal gut microbes to create butyrate, which in turn has positive effects on the gut mucosa (32, 33). L-nicotinamine is utilized in various pyridine nucleotide cycles involved in NAD or NADP salvage (PMID: 21953451) and we found that this significantly goes down with age. N-acetylneuraminate is the predominant sialic acid used in cells. Leucyl-phenylalanine can be combined with formyl-methionine to make N-formylmethionyl- leucyl-phenylalanine (FMLP) which acts as a neutrophil chemoattractant and macrophage activator (34). Lastly l-arginine is a basic amino acid, but is also used as a substrate to generate nitric oxide, which has beneficial effects for the brain and memory (35). All these metabolites have a direct effect on age related co-morbidities and stress pathways. The commensal bacteria themselves have major differences in lipid metabolism, amino acid metabolism and xenobiotic processing capabilities between old and young animals. When we looked at the metabolite-level differences in the diet subtracted fraction of the microbial metabolites (the secretome), many of the metabolites were found diminished in the older maquaques that protect mammalian cells from toxicity and modulate immune cells (phenyl phosphate, biuret, methyl chloride), protection from neurotoxicity (Thiobenzamide), modulates sphingolipid and ceramide metabolism (enol-oxaloacetate) and maintains microbial homeostasis, is anti-viral and anti-carcinogenic (4-fluoroglutamate, (2Z)-2-amino-3-hydroxy-3-(4- hydroxyphenyl)prop-2-enoate, (2S,3R)-2-[(2-aminophenyl)amino]-3-carboxy-3-oxopropanoate). All of these changes feed into stress-inducing pathways within the host as shown with serum IROA analysis before.

In this study, a plethora of bioinformatic and high-throughput methodologies were used in this study to link age-associated microbial changes and corresponding changes in microbial metabolites to age-associated immune and metabolic abnormalities. This essentially imply that microbial manipulations can prove to be an effective tool in mitigating several age-associated morbidities in the aging population.

In summary, the rhesus macaque model showed to be a reliable model for studying age-associated changes in the microbiome and immunity. The correlation and metabolite analysis has also given us a future direction for asking more specific questions about how age-associated changes in the microbiome link to age-associated changes in the immune system. Future studies will address these questions and attempt to elucidate these mechanistic links. Understanding how the microbiome and the immune system affect each other could open the door to better therapeutic interventions for immune and metabolic diseases.

## Materials and Methods

### Animals

We enrolled 12 Indian Rhesus Macaques (RM), consisting of 8 old animals (age ≥ 18 years old) and 4 young animals (age 3-6 years old. Animals were housed and blood/tissue/stool samples collected at the New Iberia Research Center (NIRC) at the University of Louisiana at Lafayette.

### Ethics Statement

All animal experiments were conducted following guidelines established by the Animal Welfare Act and the NIH for housing and care of laboratory animals and performed in accordance with institutional regulations after review and approval by the Institutional Animal Care and Usage Committees (IACUC) at the NIRC. All efforts were made to minimize suffering.

### Analysis of plasma markers of immune activation and inflammation

Plasma inflammation was determined using either an ultra-sensitive, multiplexed Milliplex magnetic bead-based assay panel (EMD Millipore) acquired on a MAGPIX instrument (Luminex Corporation) or commercially available ELISA tested on RM and analyzed using a spectrophotometer (Biotek Corportation). Multiplex assay was used in the measurement of plasma levels of IL-2, IL-6, IL-8, IL-10,, IFN-γ, and TNF-α. Commercial ELISA kits were used in the measurement of plasma levels of, hsCRP (Life Diagnostics), and Neopterin (IBL international).

### Phenotypic analysis of Immune cells subsets by Flow Cytometry

Flow cytometry was performed on peripheral blood mononuclear cells (PBMCs) from rhesus macaques using a 15 color flow cytometry panel consisting of CXCR5:FITC, CD4:PerCP-Cy5.5, IL-21R:APC, HLA-DR:AF700, Ki67:APC-Cy7, CCR7:BV421, PD1:BV605, BTLA:BV650, CD8:BV711, CD150:PE, CD95:PE-Texas Red, CD28:PE-Cy5, ICOS:PE-Cy7, CD3:BUV395, and Live/Dead viability dye in aqua channel. Briefly, PBMCs were isolated from whole blood using density centrifugation using Ficoll separation. After isolation, PBMCs were shipped via cryoshipper to the University of Miami and stored in liquid nitrogen. PBMCs were thawed and rested overnight before extracellular and intracellular staining. After staining, cells were acquired on Fortessa flow cytometer using FACS Diva software, then analyzed using Flowjo and Prism software.

### Gut microbiome analysis

Fresh fecal samples were collected from anesthetized animals by insertion of a fecal loop into the rectum and swiping several times. Fecal samples were kept in 1.5 mL microcentrifuge tube and shipped on dry ice to the University of Miami, then stored at −80°C. Microbial DNA was isolated from fecal samples using DNeasy Powersoil Kit (Qiagen) and quantified using Qubit DNA analyzer (Invitrogen/Life technologies). DNA samples were sent to the University of Minnesota Genomic Center for whole genome sequencing of the microbial profile. MetAMOS or PartekFlow softwares were used for microbial profiling.

Shotgun metagenomics library was constructed with the Nextera DNA sample preparation kit (Illumina, San Diego, CA), as per manufacturer’s specification. Barcoding indices were inserted using Nextera indexing kit (Illumina). Products were purified using Agencourt AMpure XP kit (Beckman Coulter, Brea, CA) and pooled for sequencing. Samples were sequenced using MiSeq reagent kit V2 (Illumina).

Raw sequences were sorted using assigned barcodes and cleaned up before analysis (barcodes removed and sequences above a quality score, Q≥30 taken forward for analyses)> For assembly and annotation of sequences, MetAMOS pipeline or Partek Flow software (Partek^®^ Flow^®^ Partek Inc., St. Louis, MO) were used. These softwares provide powerful tools to filter unique hits between human and mouse-specific genes versus microbial signatures. Alpha and Beta diversity calculations were done using embedded programs within the metagenomics pipeline, or using Stata15 (StataCorp LLC, College Station, TX) or EXPLICET software. Functional profiling was performed using HUMAnN2-0.11.1 (38) with Uniref50 database to implement KEGG orthologies.

### Dimensionality reduction and bacterial association analysis

We utilized OTU-matrix based dimensionality reduction and clustering algorithms. Compared to Qiime-derived PCA, which is done sample-wise, these algorithms are identity agnostic and decipher qualitative association (and disassociations) between experimental groups. We used Graphia software (Edinburgh, UK) and its implementation of Markov Clustering algorithm (MCL) (39). Using our group-wise OTU matrix, MCL looked for cluster structures using mathematical bootstrapping. While this method is agnostic to phylogenetic hierarchy in the matrix, we used the hierarchy as identifying markers to understand the bacterial clusters and changes in those clusters within the 3 study groups. MCL used the stochastic flow of the matrix to decipher the distances between the OTUs at equilibrium, thereby generating a cluster map by using correlation scores as distance. For generating the cluster, nodes scoring above a Pearson correlation value of 0.85 were used.

### Metabolite analysis by mass-spectrometry

To determine the impact of the microbiome on host metabolism in our rhesus macaque model, we used mass-spectrometry based Isotope Ratio Outlier Analysis (IROA^®^) from RM plasma and feces. This pipeline encompasses sample preparation, LC-MS based peak acquisition, proprietary software-based library creation, normalization and quantification of metabolites. IROA^®^ offers a unique platform to create and normalize a local library and account for run-to-run variability over years of acquisition using the internal standards (IROA^®^-IS) and long-term reference standards (IROA^®^-LTRS). IROA^®^ discerns peaks from noise and artifacts by spiking biological samples with labeled isotopic standards containing 95% C^13^. The combination of the naturally-occurring and U-13C 95% metabolites form a unique, reciprocal pattern of peaks during mass spectrometric analysis is classified to IROA^®^-LTRS derived library bins based on parsed ‘smiley’ peak patterns in the spectra. This pattern is used to discern veritable peaks from noise and artifacts in the sample. We have standardized and validated this method in our laboratory (24). LC-MS acquisition was done using the same instrument parameters as describes for IROA and analysis was done with MZmine2 with integrated Metacyc database.

### Statistical analysis

Data between young and old animals were compared using student’s unpaired T test. For correlation analysis, Microbial data was reordered according to difference in prevalence between young and old animals of a given microbe in the microbiome. The 25 microbes most prevalent in old or young animals was then correlated with plasma cytokines using prism software. Correlation analysis were performed using Spearman rank order correlation using prism software. Heat maps were generated showing the results and significant correlations. Results with a p value <0.05 were considered significant.

## Acknowledgments

This study has been funded by grants awarded to Savita Pahwa (R01AI123048), Laboratory Sciences Core of the Miami Center for AIDS Research (P30AI073961) and Institutional research support to Santanu Banerjee. We would like to acknowledge the University of Minnesota Genomic Center for shotgun sequencing.

---

**Supplementary Figure 1:**
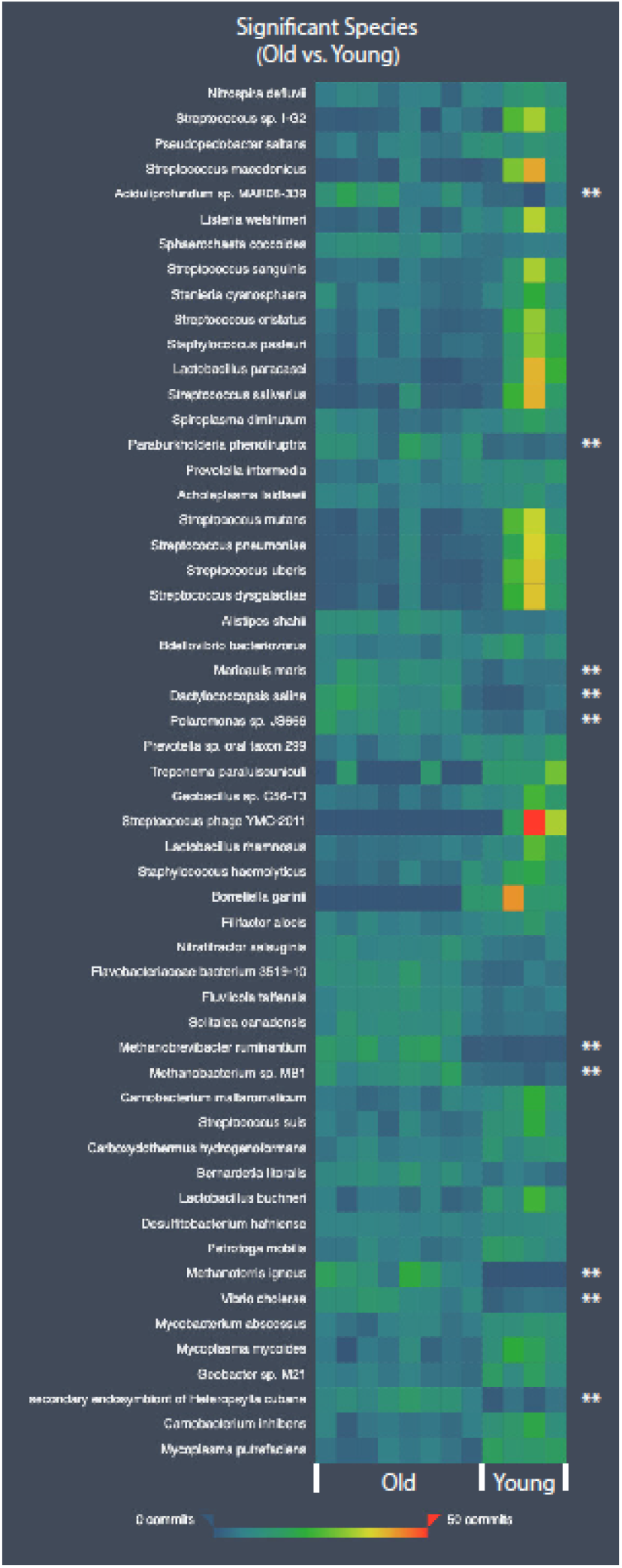
Heatmap showing species-level differences between old and young animals. All species have significant differences between groups. Species significantly higher in old animals are marked with ‘**’. Test of significance was pairwise Mann-Whitney U test and p<0.05 was considered significant

**Supplementary Figure 2:**
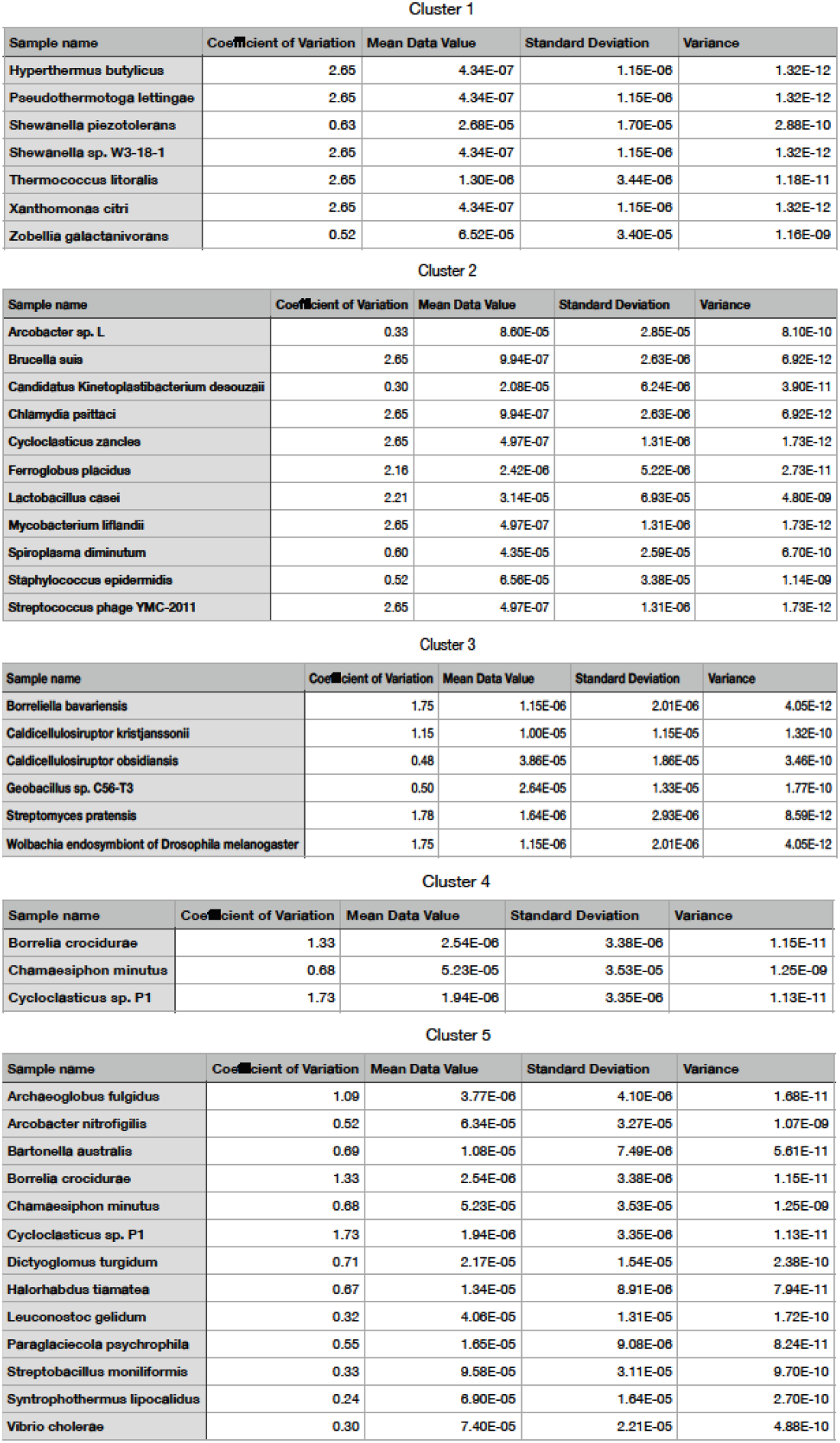
Species composition of individual microbial clusters identified for old animals in main Figure 6.

